# Tick-box exercise: Choice architecture of online classified advertisement websites can improve reporting of CITES Certificate ownership

**DOI:** 10.1101/2025.10.06.680839

**Authors:** Jon Bielby, Rebecca Graylish, Eve Power, Evangeline Button

## Abstract

The ethical and sustainable trade of nature and natural products relies on the presence of and compliance with legislation. The Convention on International Trade in Endangered Species (CITES) has been the global standard for the past half century, but doubts exist about compliance with CITES because of the lucrative nature of illegal trade, and because of misunderstandings regarding the details of specific in-country regulations. For two commonly traded species requiring UK CITES Article 10 (A10) Certificates we collected data on the frequency of reporting of the correct paperwork. We did so on two UK websites that vary in their format: one contains both a free text field and a tick box to capture information on CITES Certificates, the other contains solely a free text field. For each species the tick box captured A10 Certificate ownership in 99% of advertisements, which was significantly higher than that reported in the free text field of the same website. Further, the presence of a tick box also encouraged higher reporting of A10 ownership in the free text field, with a significant difference between websites. These results suggest that the choice architecture of websites can have a significant impact on how effectively advertisers report ownership of the correct Certificates and information.

## Introduction

The global trade in pets and ornamental plants is taxonomically diverse and economically lucrative (Hughes 2021; Watters et al. 2022), and its size has resulted in a range of concerns including the decline of wild populations due to unsustainable harvesting (Tingely et al. 2017; Hinsley et al. 2018). Legislation aimed at a more sustainable use of animals and plants is one method that aims to prevent the decline and extinction of wild populations (Sinclair et al. 2021), the most notable of which is the Convention on International Trade in Endangered Species (CITES 1973).

CITES lists species on appendices (I-III) according to the level of risk posed by trade to their long-term survival. However, within CITES processes and implementation there remains flexibility within member states, and in some circumstances regional categorisation may be higher than that of the global CITES appendices. An example of this are European tortoise species (*Testudo graeca, T. hermanni*, and *T. marginata*) within the EU and UK, where the species are listed as Annex A (the EU and UK system’s equivalent to Appendix I), a higher level of listing than their CITES global categorisation within Appendix II. On an operational level, this means that within the EU and UK it is necessary to hold a CITES Article 10 (hereafter A10) Certificate for the display or sale of these species (UK Government 2025) and that Certificate numbers should be displayed in any advertisement (Section 6, The Control of Trade in Endangered Species Regulations 2018).

As with any intervention aimed at fostering behaviour change, compliance with CITES legislation is imperfect. At the broadest scale, the illegal trade in some Appendix I species is considerable (Challender and McMillan 2014; Waeber et al. 2023), with complex trade dynamics (Challender, Harrop, McMillan 2015). At a finer scale, when focusing on a single species, in a single country, via a single route of sale, information related to a seller holding the required A10 Certificate may be poor (consciously or otherwise). For example, only 16% of the text fields in online classified advertisements of *Testudo hermanni* contained reference to the necessary A10 Certificates (Bielby et al. 2023). In the latter case there may have been a shortfall in advertiser knowledge of what an advertisement should contain to meet website guidelines, comply with existing legislation, and effectively make a sale. This poor level of communication related to the necessary certification within a legal and easily monitored route of sale suggests that conservation- and welfare-oriented interventions need to better support and direct people towards including the necessary information.

For online advertisements there are numerous ways in which the website-user interface can be designed with a view to meeting their objectives (Jameson et al. 2014). Specifically, the environment in which users make decisions on which information to include in an advertisement (the ‘choice architecture’; Münscher, Vetter, Scheuerle 2016) can incorporate features aimed at influencing the decisions that people make (e.g., the information they include). These features are often called ‘nudges’ (Thaler 2018). Typically, online classified advertisement sites have free text fields to allow an item description and related details to be listed. However, while free text fields are beneficial for expanding on details, the lack of restrictions when formulating text can cause a substantial cognitive load (Gwidzka 2010), leading to uncertainty about the most relevant information to include. Tick boxes are an aspect of choice architecture that can nudge users to include the information necessary to ensure a legal and sustainable trade, such as the presence of relevant licenses or permits. Tick boxes are often used (to varying degrees of success) in the online advertisement of age-restricted physical products such as alcohol (Williams and Ribisl 2012), and tobacco (Williams, Ribisl, Derrick 2015), pre-anaesthesia health-checks in humans (Webb and Wilson 2017), and in the verification of professional certification (for example, for healthcare practitioners purchasing restricted medical supplies; General Pharmaceutical Council 2025). Alone, or in combination with requests for further certification details (e.g., license or permit numbers), they can supplement free text fields to improve reporting of the necessary information related to website and legal guidelines. To our knowledge, however, the effects of choice architecture design within online advertisement of pets is unknown.

In this study, we investigate how information related to A10 ownership varies in free text fields compared to that in tickboxes. Specifically, for two popular classified advertisement websites and two commonly traded and kept UK species, the African Grey Parrot (*Psittacus erithacus*) and Hermann’s Tortoise (*Testudo hermanni*), we addressed the following questions:

a. Did websites vary in their level of advertisement of A10 ownership based solely on their free-text fields?
b. Did the additional presence of an A10 tickbox and ‘Certificate Number’ field improve the reporting of A10 ownership compared to the free text of both websites?

Based on previous research in the online trade of pets in the UK (Bielby et al. 2023) we predicted that free text would not frequently include information on A10 Certificates and that tick boxes would result in a higher proportion of advertisements including details of A10 Certificates (McCullough et al. 2009; Webb and Wilson 2017). Given the current level of scrutiny on online sales (Blue Cross for Pets & Born Free Foundation 2022; BVA 2023) and the trade in ‘exotic’ pets in the UK’s devolved governments (Scottish Animal Welfare Commission 2022), we hoped to identify whether the choice architecture of online classified advertisements could encourage advertisers to improve their reporting of the necessary permits and optimise compliance with website guidelines, UK legislation, and the aims and objectives of CITES.

## Methods

Two commonly used websites for animal classified advertisements were the basis for our data collection, which are referred to here as Site A and Site B. Both websites advertise pets of different types, but have notably different structure and choice architecture. These differences made the two sites ideal for comparing the efficacy of tick boxes and free text fields in recording how frequently A10 Certificate ownership was communicated in advertisements. Specifically, Site A contains a free text field, a tick-box field named “CITES Article 10”, and a related field “CITES Article 10 Certificate Number”. In contrast, Site B contains a free text field, and while it does contain a tickbox asking the advertiser “Is Endangered Species”, there is no clear reference to Annex A species or Article 10 Certificates. When posting an advertisement on Site A, it is possible to select the option “No – this species does not require an Article 10 Certificate”, regardless of the species within the advertisement being placed, thereby making it possible for users to give incorrect information.

We focused data collection on two of the most commonly traded species requiring A10s: the African Grey Parrot (*Psittacus erithacus*) and Hermann’s Tortoise (*Testudo hermanni*). Both species have experienced extreme population declines in the recent past. *Testudo hermanni* is currently listed as Vulnerable on the IUCN Red List according to criteria A2acde+4acde (Luiselli 2024), which indicates an observed, estimated, inferred or suspected decline population numbers of over 30% that is either in the past or is ongoing (IUCN Red List Cats and Criteria). *Psittacus erithacus* is currently listed at a higher level of risk, being Endangered according to criteria Endangered A2bcd+3bcd+4bcd (Birdlife 2021). Both species are listed by the IUCN Red List assessments as showing continued population decline, and the threat processes included for both species include “Biological resource use: Hunting & trapping terrestrial animals”. Accordingly, *P. erithacus* has been included on CITES Appendix I since 2017 (Species+ 2025) and *T. hermanni* on Appendix II (Species+ 2025) since 1975, and Annex A since 1997 when the EU Wildlife Trade Regulations were introduced, meaning that within the EU and UK both species are subject to CITES Article 10 certification requirements. Previous research suggests that online trade in both species has either not effectively communicated ownership of A10 Certificates or that the online trade may be used as a cover for laundering illegally sources and owned animals. Bielby et al (2023) highlighted a low level of communication of A10 ownership in *T. hermanni* advertisements on a single UK website, suggesting improvements in the system are possible. *P. erithacus* are a highly traded species that would benefit from effective monitoring and identification of individuals (Poole and Shepherd 2017; Martin, Senni, D’Cruze 2018; Martin, Senni, D’Cruze, Bruschi 2019; Ameziane et al. 2024; Davies, D’Cruze, Martin 2024). For both species there is also evidence that legal and illegal trade networks overlap (Biello et al. 2021, Davies, D’Cruze, Martin 2024).

Our data collection took place from 28^th^ February until 9^th^ July 2025. We used the terms “Hermann Tortoise” and “African Grey” as search terms in free text fields or “Parrots” from preset drop-down menus to identify suitable advertisements. For each advertisement we visually interrogated the text fields accompanying the advertisements for any language that could signify ownership of CITES Article 10 Certificates. To be as inclusive as possible we used the following search terms: “A10”, “CITES”, “license”, “paperwork”, “permit”, “microchip”, and obvious misspellings and truncations therefore. For each advertisement we therefore scored with a 1 or a 0 whether the free text field of the advertisement contained reference to ownership of the relevant paperwork and whether or not any Certificate details were included (i.e., the Certificate number). Further, for advertisements placed on Site A we recorded whether the “CITES Article 10” tick box and related “CITES Article 10 Certificate Number” fields were completed. To avoid duplicated advertisements within a website we noted the CITES permit number and/or the advertisement ID number, both of which were unique to a single advertisement. Our final data therefore consisted of advertisements for two species (*P. erithacus; T. hermanni*) on two sites, with each advert being scored for whether the free-text field contained reference to ownership of a CITES A10 Certificate, and for each advert on Site A whether a tickbox had been correctly completed. From these data we calculated the proportion of advertisements communicating ownership of an A10 Certificate for each combination of website, species, and feature of choice architecture (free text or tick box).

To identify how species, website, and choice architecture affected the likelihood of communication of A10 Certificate ownership for each combination we compared proportion of advertisements communicating A10 ownership with a two proportion z-test using the prop.text function in base package of the software R (R Core team 2025). Specifically, for each of our two focal species we compared the proportion of advertisements communicating CITES A10 Certificates between choice architecture features within one website (Site A), and between the text fields of the two sites. This yielded four comparisons in total:

*Testudo hermanni*, Site A tick box Vs Site A free text;

*Testudo hermanni*, Site A free text Vs Site B free text;

*Psittacus erithacus*, Site A tick box Vs Site A free text;

*Psittacus erithacus*, Site A free text Vs Site B free text;

## Results

### African grey parrot (*Psittacus erithacus*)

Within site comparison in the African Grey Parrot highlighted that a tickbox (60/61) was significantly more effective than a free text field (41/61) in capturing the ownership of CITES A10 Certificate (p<0.001). Similarly, free text fields in Site A (41/61) and Site B (26/71) exhibited significant variation in their inclusion of the correct Certificate information (p<0.001).

**Figure 1.**
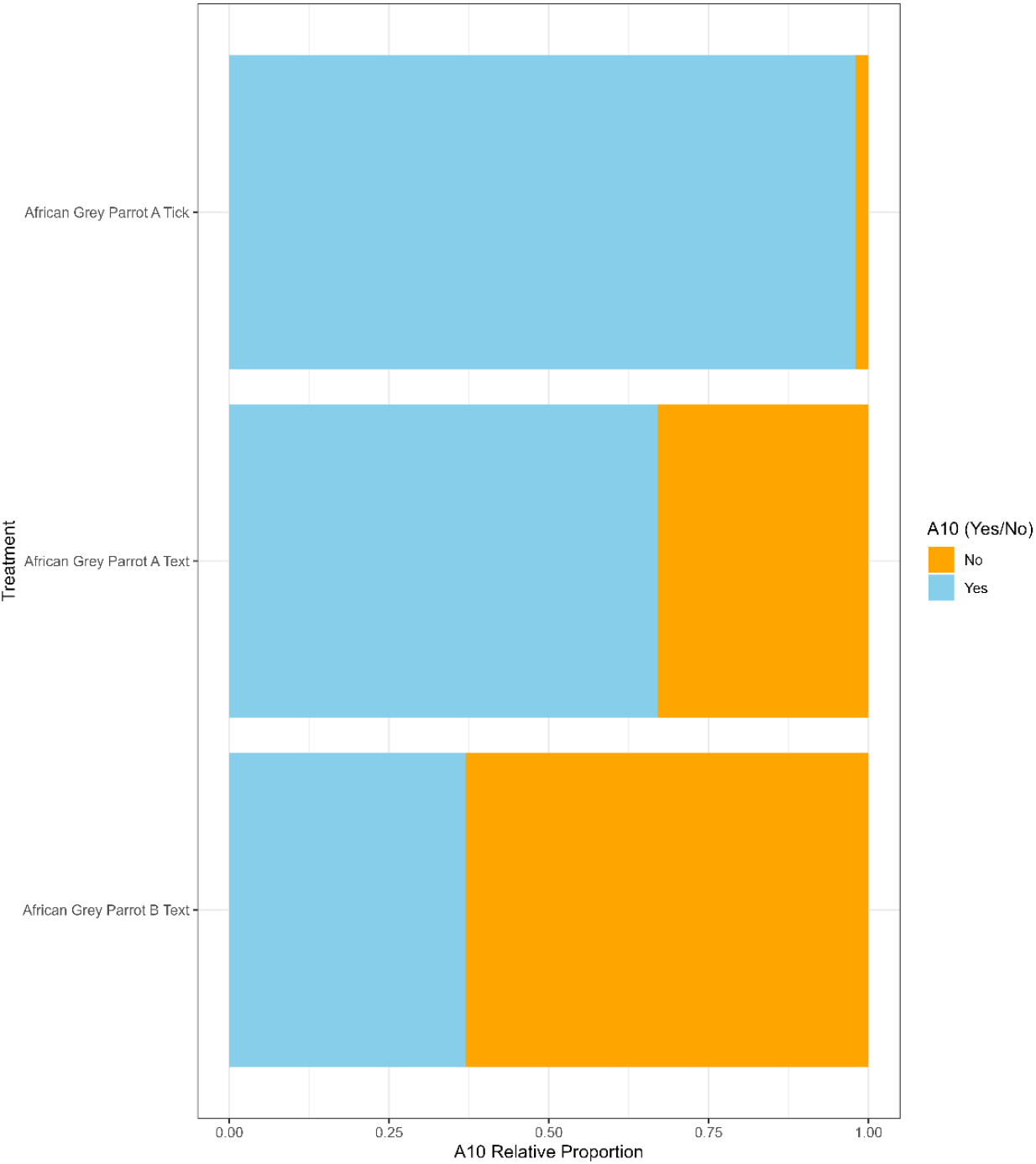
Proportion of total advertisements of African Grey Parrots (*Psittacus erithacus*) communicating ownership of CITES Article 10 Certificates for each of Site A tick box (n=60/61), Site A free text (n=41/61), and Site B free text (n=26/71). Two proportion z-tests indicated that proportions of A10 details in Site A text and tickboxes significantly varied from each other, as did Site A and Site B free text.

### Hermann’s tortoise (*Testudo hermanni*)

Within site comparison in Hermann’s tortoises suggest that a tickbox (64/65) was significantly better than a free text field (50/65) in communicating the presence of a CITES A10 Certificate (p<0.001). Between site comparison of free text fields outlined a similar difference with Site A (50/65) and Site B (48/102) exhibiting significant variation in their inclusion of the correct Certificate information (p<0.001).

**Figure 2.**
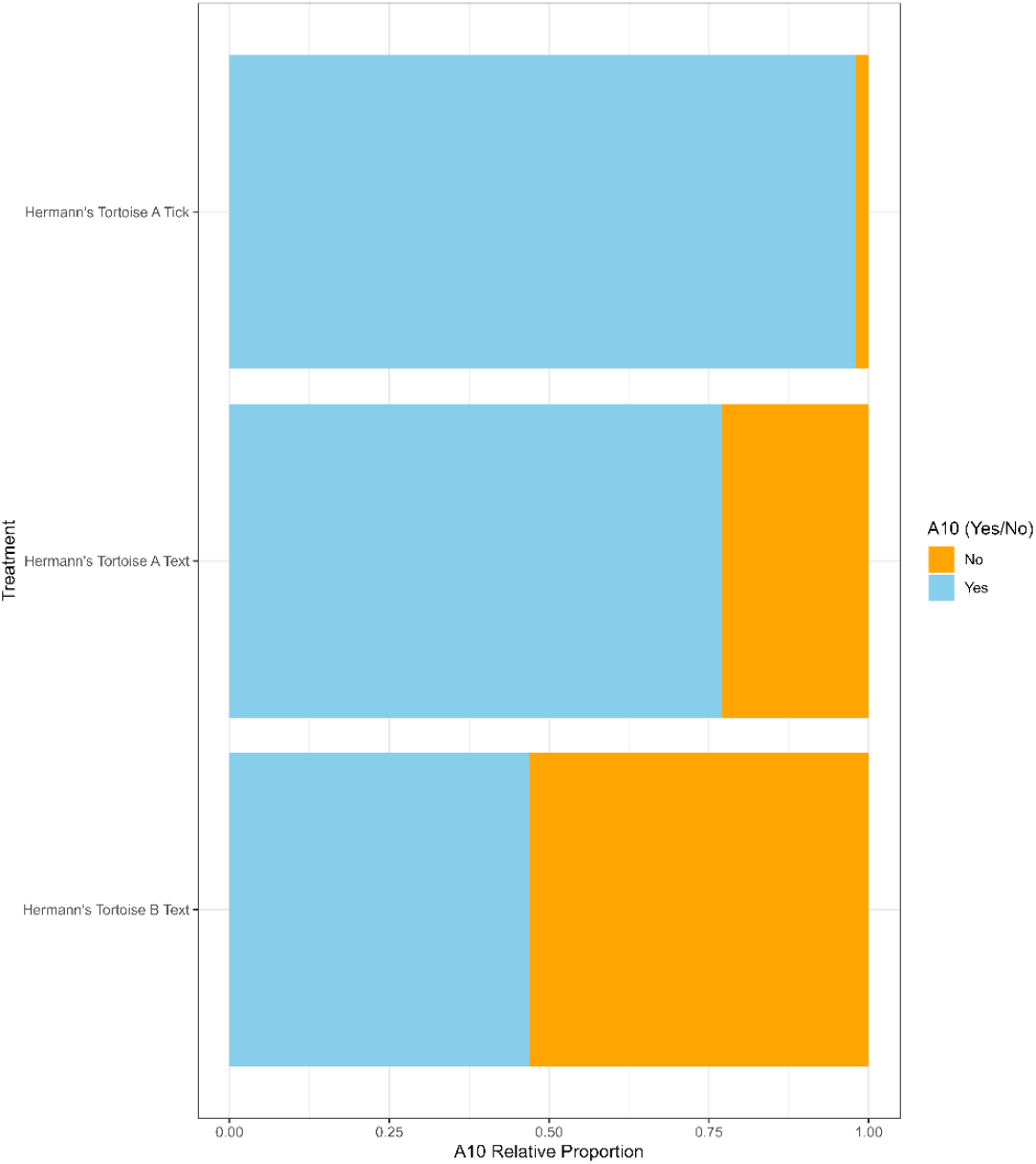
Proportion of total advertisements of Hermann’s Tortoise (*Testudo hermanni*) communicating ownership of CITES Article 10 Certificates for each of Site A tick box (n=64/65), Site A free text (n=50/65), and Site B free text (n=48/102). Two proportion z-tests indicated that proportions of A10 details in Site A text and tickboxes significantly varied from each other, as did Site A and Site B free text.

## Discussion

The size and scope of the trade in pet animals and ornamental plants has directly led to the decline and local extinction of multiple species (Marsh et al. 2022; Challender et al. 2023). Effective legislation to prevent overharvesting while promoting sustainable use of natural resources is important for conserving biodiversity and meeting obligations to the UN Sustainable Development Goals (UN DESA 2024). CITES appendices have been used for almost half a century with this in mind, but there are concerns about the level of compliance within this existing framework. In part, lack of compliance may be due to purposely illegal activities (Wyatt et al. 2020), but in some circumstances it may be a result of lack of clarity around regulations (Rudolph and Riley 2017), in this case details of what paperwork and information is required when buying or selling animals and plants. The data presented in this study suggest that within the UK the choice architecture of classified online advertisement websites can significantly affect the level of reporting ownership of the necessary Certificates and permits. Specifically, the addition of a simple tick box and a short text field for reporting the A10 Certificate number significantly increased the reporting of CITES Certificates in two widely traded species: Hermann’s tortoise and the African grey parrot. The broader implications are that some simple interventions improve transparency within the UK’s legal trade in exotic pets, the importance of which is only likely to increase given current policy discussions surrounding the future of the trade and hobby (Born Free & RSPCA 2021; BVA 2023; SSPCA 2025).

To our knowledge this study represents the first focused effort to investigate the effect of choice architecture on aspects of the legal online pet trade. However, in other fields such as human health care there is a better evidence base evaluating the efficacy and informing the use of features such as tick boxes (Williams and Ribisl 2012; Williams et al. 2015; Webb and Wilson 2017). These studies suggest that while the use of tick boxes can improve desired outcomes, they do not provide a silver-bullet solution to meeting best practice guidelines or the law, and they must be used with caution and monitored appropriately. Their limited efficacy as a nudge to website users can be broadly attributed to two aspects of human behaviour – first, people sometimes forget things; and second, people sometimes want to cheat. For example, research into the sale of age-restricted products (alcohol and tobacco) to minors in the US suggests that tick boxes alone did not effectively reduce the illegal purchase of these products (Williams and Ribisl 2012; Williams et al. 2015; Webb and Wilson 2017). Even when the websites in question required age-verification via a driver’s license at point of purchase, the purchase could still be made if the participant was allowed to cheat (by entering a parent’s license number). However, making it harder to cheat by including stricter age-checks (e.g., by requesting social security numbers) or removing a participant’s ability to use a parent’s license number did result in better compliance (i.e., sales were refused). Combined, these results suggest that if consumers are actively trying to make an illegal purchase, then tickboxes and requests for permit details would need to be non-cheatable to effectively ensure compliance. In the context of our study this would mean a specific format of Certificate number, or greater traceability of A10 Certificate numbers and the animals to which they refer. Aside from those who actively aim to cheat the system, the use of tick boxes can be an effective, if not foolproof, way to improve compliance with best practice guidelines. In these cases they may play the role of a memory aid to prompt a participant’s memory of what information is relevant to the task in hand. For example, inclusion of tick box questions has significantly improved outcomes when added to medical surveys as a reminder to request specific information about a patient’s smoking history and plans for quitting (McCullough et al. 2009; Webb and Wilson 2017). However, despite improved outcomes in these cases the use of tick boxes could not be relied on as a sole method of improvement, as efficacy was not 100%.

In our data CITES A10 Certificate ownership was captured by tickboxes in 98% of advertisements (124/126) across the two species. This suggests that although not perfect, the addition of this choice architecture could improve compliance with industry standards and best practices (e.g., PAAG 2025) by both individuals placing advertisements, and the websites upon with they were placed. Further, the inclusion of tickboxes also increased reporting of A10 Certificates within associated free text fields (i.e., for both species the text field reporting of A10 ownership was significantly higher on Site A, which included a tickbox). This point speaks to the importance of website features both for directly capturing relevant information and also as a nudge in other aspects of website design. To better understand the necessity of nudges it would be useful to quantify the awareness of CITES paperwork requirements and the reasons behind them for both sellers and buyers, both online and more generally. This would allow a more detailed understanding of when and where choice architecture is likely to be effective, and where more focussed information would provide a better foundation for improving compliance.

The implications of these findings could extend beyond A10 certification within UK classified advertisement sites. The subject of keeping exotic pets is currently being discussed within the UK’s devolved governments and new policies and legislation could be implemented in the future (Scottish Animal Welfare Commission 2022; BVA 2023). Some legislation that has gained traction among advocates of change in the exotic pet sector involve the use of ‘Positive lists’, which are a list of species that may be legally kept (Toland et al. 2020), and pre-purchase knowledge checks (BVA 2023; Loeb 2025). For either or both suggested interventions to be effective, one important aspect would be the ability to verify the species identity of the animals being traded, and the certification of the owner’s knowledge status. Our analyses suggest that both aims would require careful thought into choice architecture and nudges required to effectively and accurately capture the necessary information when animals and/or plants are traded, whether online or in person. Clearly, though, there are major limitations to this approach and in many cases simple tickboxes and other similar behavioural nudges would be insufficient in efforts to ensure compliance. A large proportion of the wildlife trade is illicit (Wyatt et al. 2020) and most points along the supply-chain are poorly known (Sinclair et al. 2021) and likely highly difficult, expensive, and dangerous to monitor and control (Wyatt et al 2020; Gore et al. 2023). The implications of this study are therefore small in scale, focussing on a relatively transparent, easily monitored route of advertisement and sale, and should not be used to advocate the use of such approaches in other, broader aspects of the pet trade. Further, even at the scale of the UK exotic pet trade it would be much more difficult to monitor and improve the efficacy of targeted legislation on less easily accessible, less transparent internet platforms (e.g., social media, dedicated forums, chat groups, the dark web; Stringham *et al*. 2021). This lack of control via these other routes of sale perhaps highlights the benefits of trying to regulate online trade effectively on engaged classified advertisement sites to avoid displacing trade to less easily monitored routes of sale. The legal trade is also commonly closely linked to illicit trade, with the mechanisms and processes of the former sometimes being used as a cover for illegal trade (Gore *et al*. 2023), which may be a particular concern in the two focal species within these analyses (Ameziane *et al*. 2024; Biello *et al*. 2021; Boratto and Griffis 2024; Davies *et al*. 2024; Mozer *et al*. 2025). Our study was only able to identify which advertisements communicated ownership of CITES Article 10 Certificates, which neither tells us whether advertisers actually possess those Certificates, nor whether the Certificates advertised were genuine. Further research to trace CITES Article 10 Certificates in a range of species could highlight useful aspects of trade dynamics such as the average number of transactions experienced by different species, and the level of actual compliance with the current regulations.

The exotic pet trade in the UK is at a crossroads. Its taxonomic diversity and size have resulted in significant welfare concerns (BVA 2023; Scottish Animal Welfare Committee 2022), leading to multiple stakeholders calling for legislative change (Elwin et al. 2020; Born Free & RSPCA 2021; SSPCA 2025). Several such changes have been advocated with a view to improving the sustainability of the trade and the welfare of the animals involved (Scottish Animal Welfare Committee 2022; BVA 2023; Loeb 2025). However, when designing and introducing interventions aimed at behaviour change it is important to carefully consider the likely compliance levels (Fairbrass et al. 2016) and how the system supports people throughout the supply chain in adhering to regulations and communicating effectively that they are doing so (Rudolph and Riley 2017). Our data suggest that some highly visible routes of trade are amenable to simple improvements that can significantly increase how well an advertiser reports their compliance with existing best practice guidelines and regulations. By extension, future interventions could benefit from coordinated efforts to design choice architecture that will support transparency and traceability throughout the supply chain and in people’s homes. Ultimately, doing so would be to the benefit of the welfare of the animals involved and the sustainability of their wild populations.

